# An AI-Native Biofoundry for Autonomous Enzyme Engineering: Integrating Active Learning with Automated Experimentation

**DOI:** 10.64898/2026.02.01.703093

**Authors:** Chuwen Zhang, Lixiang Yang, Yanjia Qin, Danjing Li, Shimao Dong, Meng Yang

## Abstract

The engineering of enzymes with novel functions is a cornerstone of synthetic biology but remains bottlenecked by the fragmentation between computational design and physical execution. While “self-driving” laboratories promise to resolve this, existing systems often rely on rigid, device-specific scripts that lack the flexibility to handle complex, evolving scientific tasks. Here, we report an AI-native autonomous biofoundry that fundamentally redefines laboratory automation through a “cloud-edge synergistic” architecture. The platform features an Agent-Native control system powered by Large Language Models (LLMs) and the Model Context Protocol (MCP), which bridges the semantic gap between abstract scientific intent and heterogeneous hardware execution. This architecture enables non-experts to orchestrate the entire Design-Build-Test-Learn (DBTL) cycle via natural language. By integrating deep phylogenetic mining, zero-shot protein language models (ESM-2), and supervised active learning, our system efficiently navigates rugged fitness landscapes. As a rigorous proof of concept, we applied this platform to evolve a Family B DNA polymerase for CoolMPS sequencing, a task requiring the incorporation of non-natural 3’-blocked nucleotides. In just three autonomous rounds, the platform achieved a hit rate of >66% and identified variants with a 37% reduction in sequencing error rate compared to a commercial reference. This work demonstrates that AI-native infrastructures can not only accelerate trait evolution by orders of magnitude but also provide a scalable, brand-agnostic paradigm for the future of automated scientific discovery.

## 1. Introduction

Enzymes function as the molecular engines of the bio-economy, underpinning critical advancements in sustainable catalysis, synthetic biology, and precision medicine [1–3]. However, naturally occurring enzymes often lack the requisite stability, activity, or substrate specificity demanded by industrial applications. For decades, directed evolution (DE) has served as the gold standard for engineering these traits [4]. By mimicking natural selection through iterative cycles of mutagenesis and screening, DE has successfully delivered biocatalysts for diverse sectors [5]. Yet, despite its Nobel-winning pedigree, traditional DE faces fundamental bottlenecks: it is inherently stochastic, labor-intensive, and resource-heavy [6]. The “you get what you screen for” paradigm often necessitates the evaluation of massive libraries (high-N) — ranging from thousands to millions of variants—to identify a single improved mutant [7]. Furthermore, navigating the vast, rugged protein fitness landscape typically relies on expert intuition or rational design, which is constrained by our incomplete understanding of complex structure-function relationships [8, 9]. This reliance on specialized expertise creates a significant barrier to entry, limiting the scalability of enzyme engineering [10].

The convergence of artificial intelligence (AI) and laboratory automation is currently reshaping this landscape, giving rise to “self-driving laboratories” [11 – 13]. On the computational front, deep learning models, particularly Protein Language Models (PLMs) like ESM-2, have demonstrated a remarkable ability to capture evolutionary patterns and predict functional hotspots from sequence data alone (zero-shot prediction) [14, 15]. Simultaneously, automated biofoundries have enabled the high-throughput execution of complex “Design-Build-Test” workflows with high precision and reproducibility [16].

However, a critical gap remains in the seamless orchestration of these technologies. Current workflows often treat computation and experimentation as isolated silos, creating “automation silos” where human researchers must manually transfer data, retrain models, and redesign libraries between rounds [17 – 19]. Moreover, most existing automated systems rely on rigid, pre-programmed scripts that lack the flexibility to handle dynamic scientific tasks or adapt to heterogeneous hardware. While recent efforts have integrated specific models with liquid handlers [17, 18], there remains a lack of a generalized, AI-native architecture that can fundamentally bridge the semantic gap between abstract scientific intent and physical execution — allowing non-experts to efficiently navigate evolutionary paths using minimal experimental data (Low-N) [20, 21].

Here, we report a generalized AI-Native Biofoundry that redefines laboratory automation through a “cloud-edge synergistic” architecture. Unlike traditional systems, our platform features an Agent-Native control system powered by Large Language Models (LLMs) and the Model Context Protocol (MCP) [22]. This architecture enables an AI agent to directly “perceive” instrument states and “actuate” physical workflows, translating natural language requests (e.g., “optimize polymerase activity”) into executable protocols without human intervention. We employ a “computational funnel” strategy: prioritizing candidates via global phylogenetic mining, jumpstarting evolution with zero-shot PLM predictions, and iteratively refining the search space using supervised active learning [23].

As a rigorous proof-of-concept, we applied this platform to a highly challenging problem: evolving a Family B DNA polymerase for CoolMPS sequencing [24]. This application requires the enzyme to efficiently incorporate unlabeled 3’-blocked reversible terminators — a bulky, non-natural substrate that wild-type enzymes poorly accept [25]. In just three rounds of closed-loop evolution, our platform autonomously identified variants with a 37% reduction in sequencing error rate and superior processivity compared to commercial standards. This work demonstrates that AI-native infrastructures can accelerate trait evolution by orders of magnitude, providing a scalable and brand-agnostic paradigm for the autonomous development of next-generation biocatalysts.

## 1. Results

### 2.1 Architecture of the AI-Native Automated Biofoundry

#### System Overview

To bridge the critical gap between computational design and physical execution, we established an AI-native automated laboratory execution management system (Fig. 1). Unlike traditional automation that relies on rigid, device-specific scripts, this platform functions as a “cloud-edge synergistic” distributed architecture, transforming the laboratory into a logically unified, programmable digital entity, a concept extending recent advances in reconfigurable workflow management [49].

**Figure 1.**
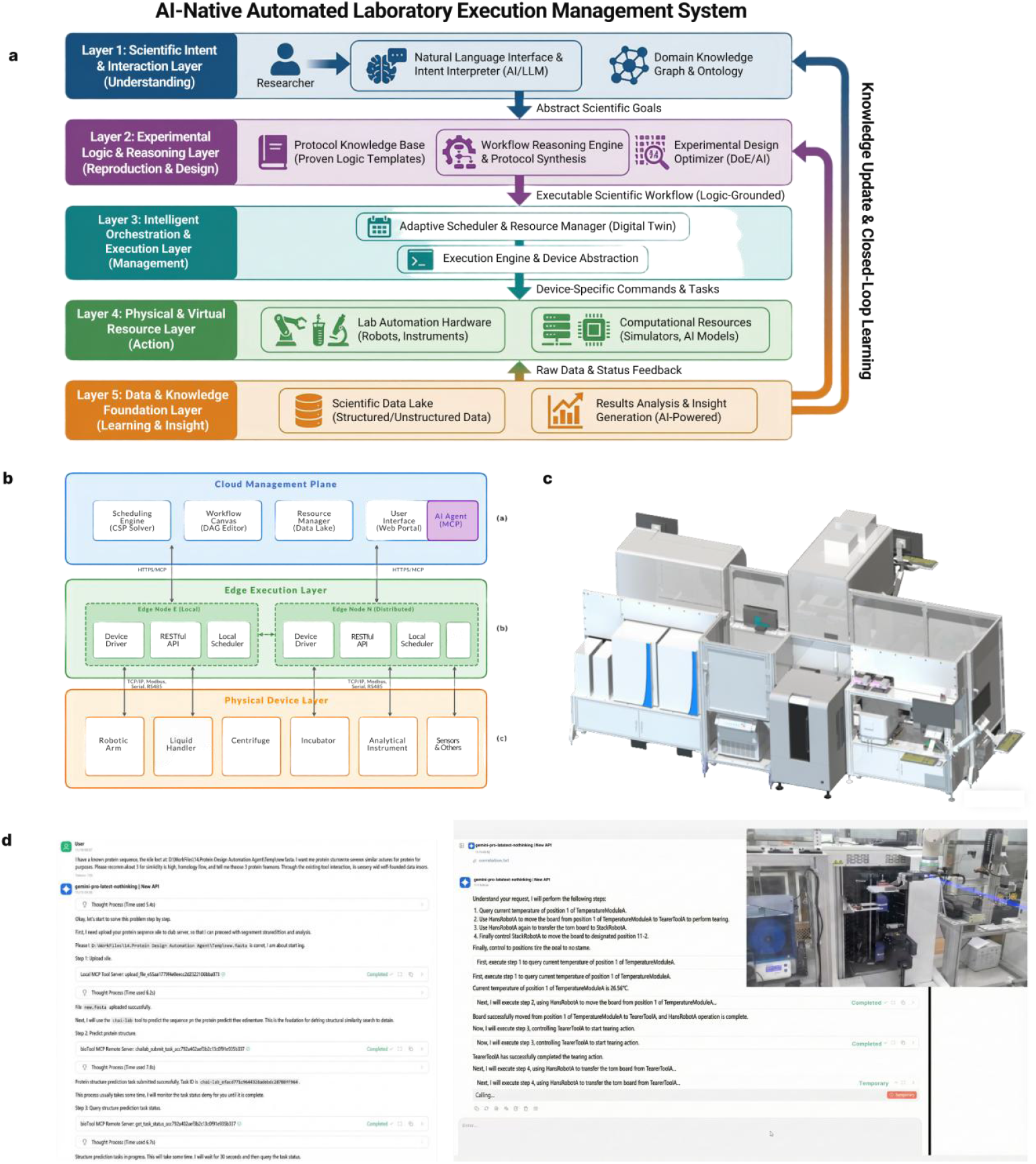
Architectural Framework and Operational Workflow of the AI-Native Automated Laboratory Execution Management System. (a) Hierarchical Functional Model: This panel illustrates the five-layer governance structure, ranging from top-level scientific intent interpretation (Layer 1) and experimental logic synthesis (Layer 2) to intelligent orchestration (Layer 3), physical/virtual resource execution (Layer 4), and the foundational data-driven knowledge closed-loop (Layer 5). (b) Cloud-Edge-Device Distributed Architecture: A detailed schematic of the “cloud-edge synergistic” design. The Cloud Management Plane handles global scheduling and digital twin synchronization; the Edge Execution Layer provides localized, high-reliability control through hardware abstraction (RESTful/MCP protocols); and the Physical Device Layer integrates heterogeneous instruments (e.g., robotic arms, liquid handlers) into a unified logical entity. (c) Physical System Implementation: A three-dimensional visualization of the reconfigurable laboratory modules, demonstrating the “software-defined topology” that allows for flexible physical layouts and rapid instrument integration. (d) Agent-Native Interaction and Analytical Output: Evidence of the system’s human-AI collaborative capabilities. It shows the transformation of natural language scientific instructions into structured, executable protocols via LLM-driven intent analysis, alongside the automated generation of experimental reports and data insights.

#### Hierarchical Governance and Cloud-Edge Synergy

The system is built upon a five-layer governance model that decouples high-level scientific intent from low-level hardware execution.

#### Cloud Management Plane

This layer handles global resource orchestration, data persistence, and user interaction. It hosts the “Digital Twin” of the laboratory, enabling “rehearsal” simulations of experimental protocols to identify collision risks and estimate execution timelines before physical actuation [50].

#### Edge Execution Layer

To ensure reliability and low latency, execution logic is deployed to local edge nodes. This architecture guarantees that high-frequency instrument control remains stable even during network fluctuations, preventing the “automation silos” often seen in centralized systems.

#### Agent-Native Integration via Model Context Protocol (MCP)

A distinct breakthrough of our system is its Agent-Native design. By deeply integrating the Model Context Protocol (MCP) [51], we bridged the semantic gap between Large Language Models (LLMs) and physical hardware. The AI Agent (powered by Qwen3) serves as the central orchestrator, capable of directly “perceiving” instrument states and “actuating” physical workflows. Instead of relying on human-written scripts, the Agent parses natural language instructions (e.g., “optimize polymerase for thermostability”) into structured JSON Task Graphs. It automatically resolves complex resource competition and scheduling constraints, enabling a truly autonomous “human-in-the-loop” free operational mode, aligning with the vision of global autonomous scientific agents [52].

#### Hardware Abstraction for Zero-Intrusive Integration

To address the challenge of hardware heterogeneity, we implemented a “Meta-Action” protocol model. Device capabilities are abstracted into brand-agnostic interfaces—such as generic Transfer or Incubate commands — rather than vendor-specific code, similar to the hardware-agnostic approaches pioneered by PyLabRobot [53]. This “Software-Defined Topology” allows for the zero-intrusive integration of heterogeneous instruments (e.g., from liquid handlers to analytical detectors). New devices can be integrated via dynamic runtime loading without system downtime, creating a flexible, plug-and-play ecosystem for scalable scientific discovery.

### 2.2 Physics-Informed AI Mining for Thermostable Polymerase Candidates

To identify robust polymerase scaffolds capable of supporting CoolMPS chemistry, we employed a “computational funnel” strategy to mine the global sequence space. Starting from the InterPro database, we filtered Family B DNA polymerase sequences for Thermophilic Archaea origin and applied SoluProt and molecular dynamics (MD) simulations to predict solubility and substrate binding stability. This rigorous in silico campaign narrowed the vast sequence space down to 160 high-confidence candidates for experimental validation.Biofoundry-Enabled Screening and Candidate IdentificationTo explore the functional diversity of these mined sequences, the 160 candidates were expressed and purified using our automated biofoundry. From this computationally selected library, 7 variants exhibited detectable nucleotide incorporation activity, validating the efficacy of our physics-informed filtering. We subsequently performed a comprehensive kinetic characterization of these active variants alongside the reference enzyme (REF) across four natural substrates (dATP, dTTP, dGTP, and dCTP).Evaluation of Substrate Incorporation EfficiencyWe first assessed the absolute incorporation activities of the discovered variants (Fig. 2d). The reference enzyme (REF) demonstrated a characteristic bias, with a strong preference for dATP (0.52 a.u.) and dTTP, consistent with the typical behavior of this polymerase family. Among the mined variants, A0A2Z2MPY8 and I3ZT69 emerged as the top performers, retaining substantial catalytic activity. Notably, while REF exhibited the highest peak activity for adenine, variant A0A2Z2MPY8 displayed a significantly more balanced substrate profile across all four nucleotides.To explicitly quantify the functional divergence, we calculated the differential activity. This comparative analysis revealed two key insights:Activity Retention vs. Trade-off: While most variants showed a reduction in dATP and dTTP incorporation relative to the highly optimized REF, several variants maintained activity levels within a functional range, proving they are robust scaffolds.Expanded Substrate Tolerance: Strikingly, variant A0A2Z2MPY8 exhibited a positive performance gain in handling G/C content. Specifically, it outperformed the REF sequence in both dGTP and dCTP incorporation. This suggests that A0A2Z2MPY8 has naturally evolved a broader substrate tolerance or reduced bias against G/C nucleotides compared to the wild-type scaffold, a trait highly desirable for sequencing applications where uniform incorporation is critical.Sequence-Function Relationship and Clustering AnalysisTo investigate whether the observed functional differences stem from global sequence divergence or specific motif variations, we performed a dual-heatmap analysis combining genotypic and phenotypic data.Genotypic Diversity: The Sequence Identity Heatmap (Fig. 2b) illustrates the pairwise amino acid homology among the characterized enzymes. The matrix reveals a diverse genetic background with identity scores spanning a wide range, confirming that our mining strategy successfully captured distinct evolutionary branches rather than minor variants of a single lineage.Phenotypic Clustering: Furthermore, we employed hierarchical clustering based on enzymatic activity profiles. This functional clustering revealed patterns that do not strictly mirror sequence homology. As shown in the dendrogram (Fig. 2d), the variants segregated into distinct functional groups. One group, led by A0A2Z2MPY8, clustered distinctly from low-activity variants, highlighting its unique “generalist” behavior—characterized by consistent activity across multiple substrates.This discrepancy between genotypic and phenotypic grouping underscores the plasticity of the active site, where moderate sequence variations can lead to drastically different substrate specificities. In summary, our Phase I screening identified novel variants that not only retain catalytic function but, in the case of A0A2Z2MPY8, offer superior performance in incorporating dGTP and dCTP, providing a valuable and balanced starting point for further directed evolution.

**Figure 2.**
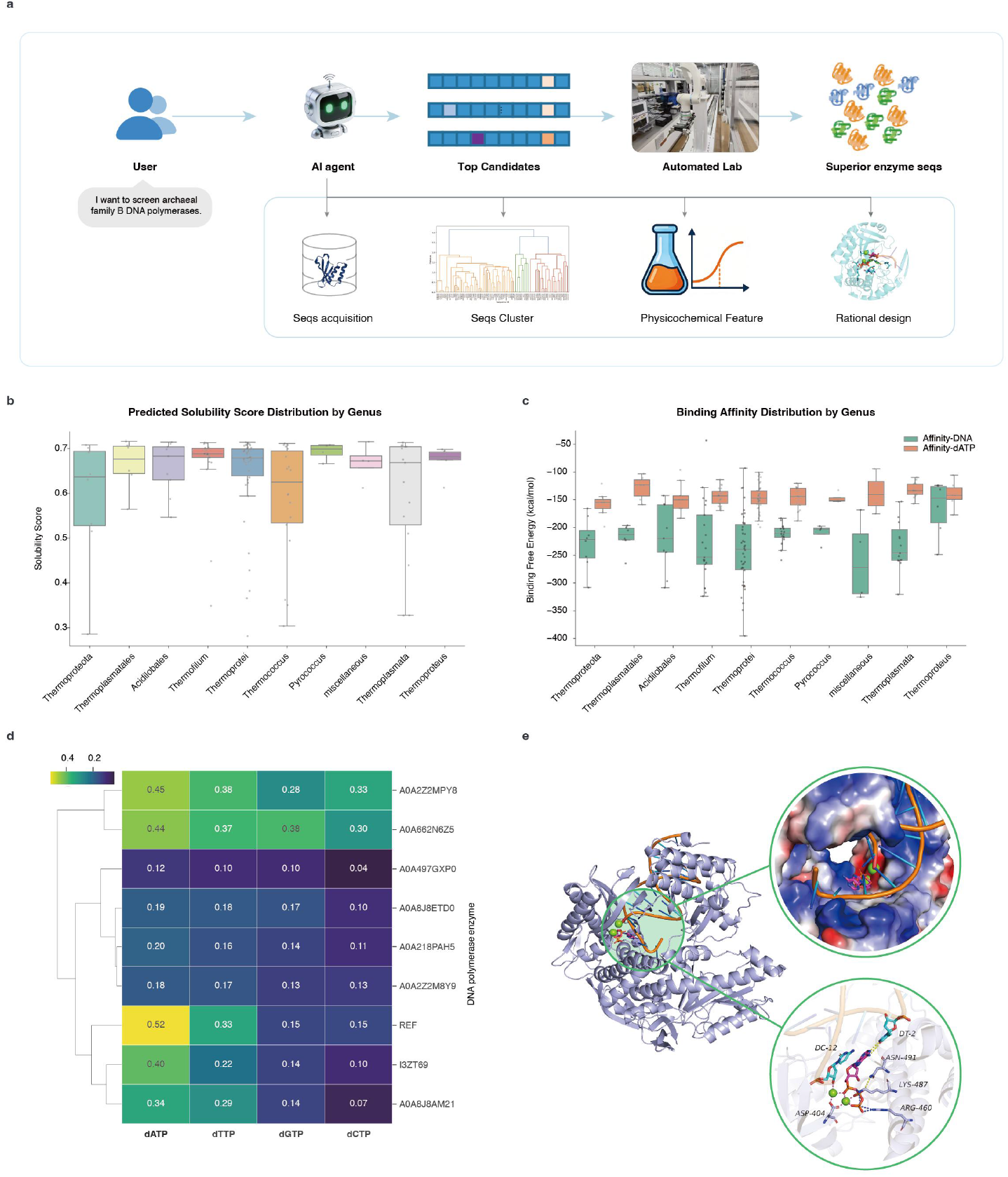
A computational-experimental framework for the discovery of novel thermostable DNA polymerases. (a) Schematic overview of the AI-driven mining pipeline. The workflow integrates database mining, phylogenetic filtering, sequence clustering, and multi-objective AI prediction to identify high-potential candidates from the InterPro database. (b) Distribution of predicted solubility scores (SoluProt) across identified thermophilic archaeal genera. Higher scores indicate a higher probability of soluble expression. (c) Molecular Dynamics (MD)-based binding free energy calculations for the top candidates. Box plots show the distribution of binding affinities for the enzyme-DNA complex (green) and enzyme-dATP complex (orange) across different genera. (d) Heatmap profiling the substrate specificity or incorporation activity (1min) of the final candidates against different nucleotides (dATP, dTTP, dGTP, dCTP), highlighting diverse activity profiles. (e) Structural validation of a representative candidate modeled by Chai-lab. The zoom-in views depict the stable coordination of the DNA template (top circle) and the incoming nucleotide (bottom circle) within the active site after 200 ns MD simulation.

### 2.3 Accelerating Evolutionary Trajectories via Active Learning

To navigate the expansive fitness landscape of the polymerase, we implemented a closed-loop engineering strategy that transitions from broad sequence exploration to focused active learning (Fig. 3). This biphasic approach allows for the efficient identification of functional hotspots followed by the precise recombination of synergistic mutations (Fig. 4).

**Figure 3.**
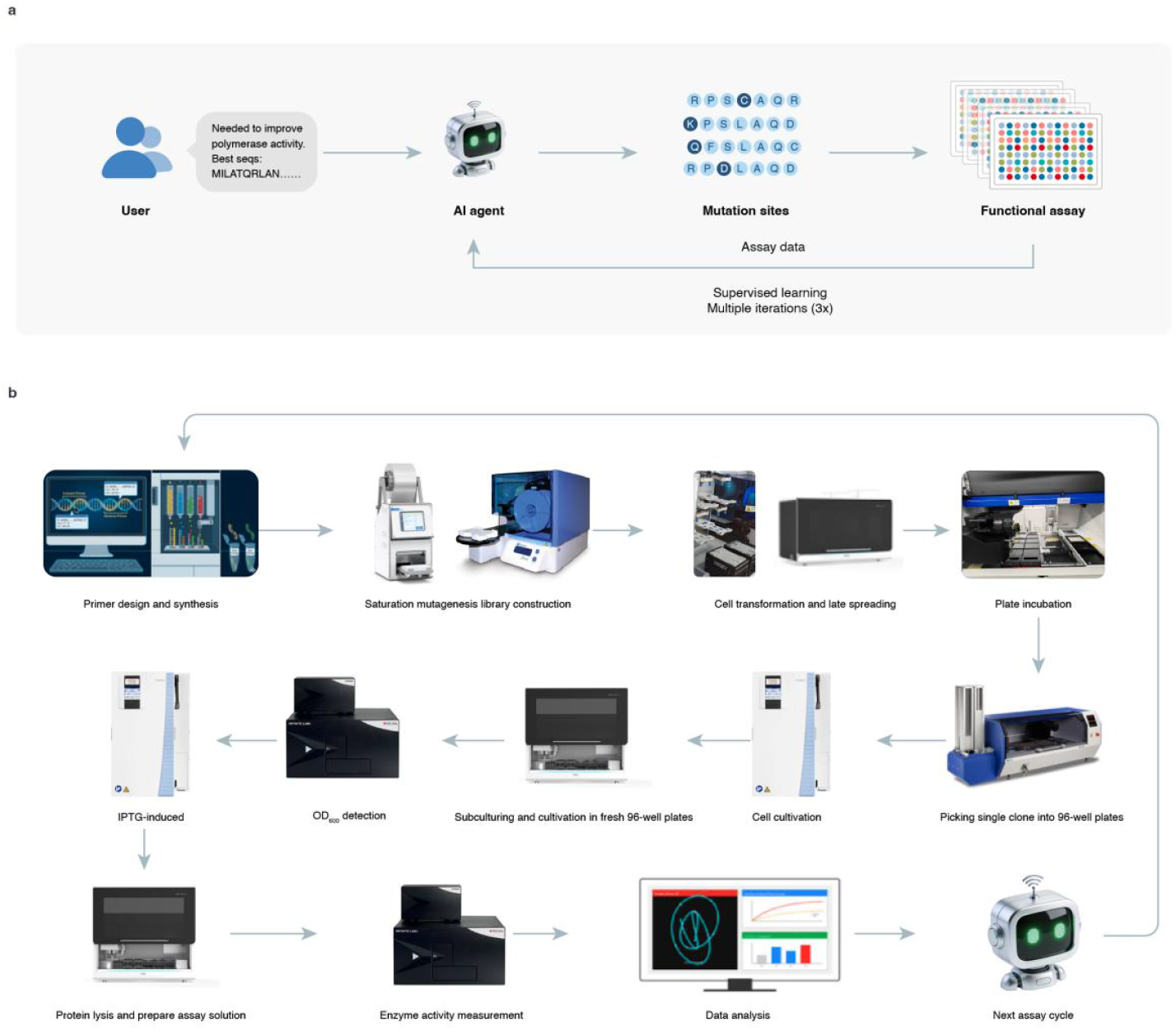
General overview of the automated protein engineering platform. a. Core modules and closed-loop workflow. The platform integrates three key modules to enable autonomous protein evolution. The process initiates with User Interaction and Initial Prediction, where users define optimization goals (e.g., improving HMT activity) via a natural language interface, and an unsupervised model (e.g., ESM-2) predicts mutation sites to generate an initial variant library. This is followed by Automated Build & Test, where an integrated biofoundry utilizes robotic systems to execute DNA assembly and functional validation. The resulting assay data drives Machine Learning-Assisted Evolution (Learn), training supervised models to predict variants with higher fitness. The system operates through iterative Design-Build-Test-Learn (DBTL) cycles, significantly reducing reliance on human domain expertise. b. Details of the BAL automated laboratory pipeline. This panel illustrates the hardware infrastructure and modular workflows. The process begins with Primer Design and Synthesis, automatically generating mutagenesis primers based on AI-identified sites. Library Construction employs modular site-directed mutagenesis (SDM) via PCR and HiFi assembly. Cell Transformation and Culture are managed by automation systems performing transformation, plating, colony picking, and incubation at the 96-well scale. Protein Expression and Lysis involve IPTG induction and crude lysate extraction. Finally, Activity Detection and Data Analysis utilize microplate readers to measure kinetic parameters, with data processed via automated scripts and fed back to the AI agent to trigger the next evolutionary cycle.

**Figure 4.**
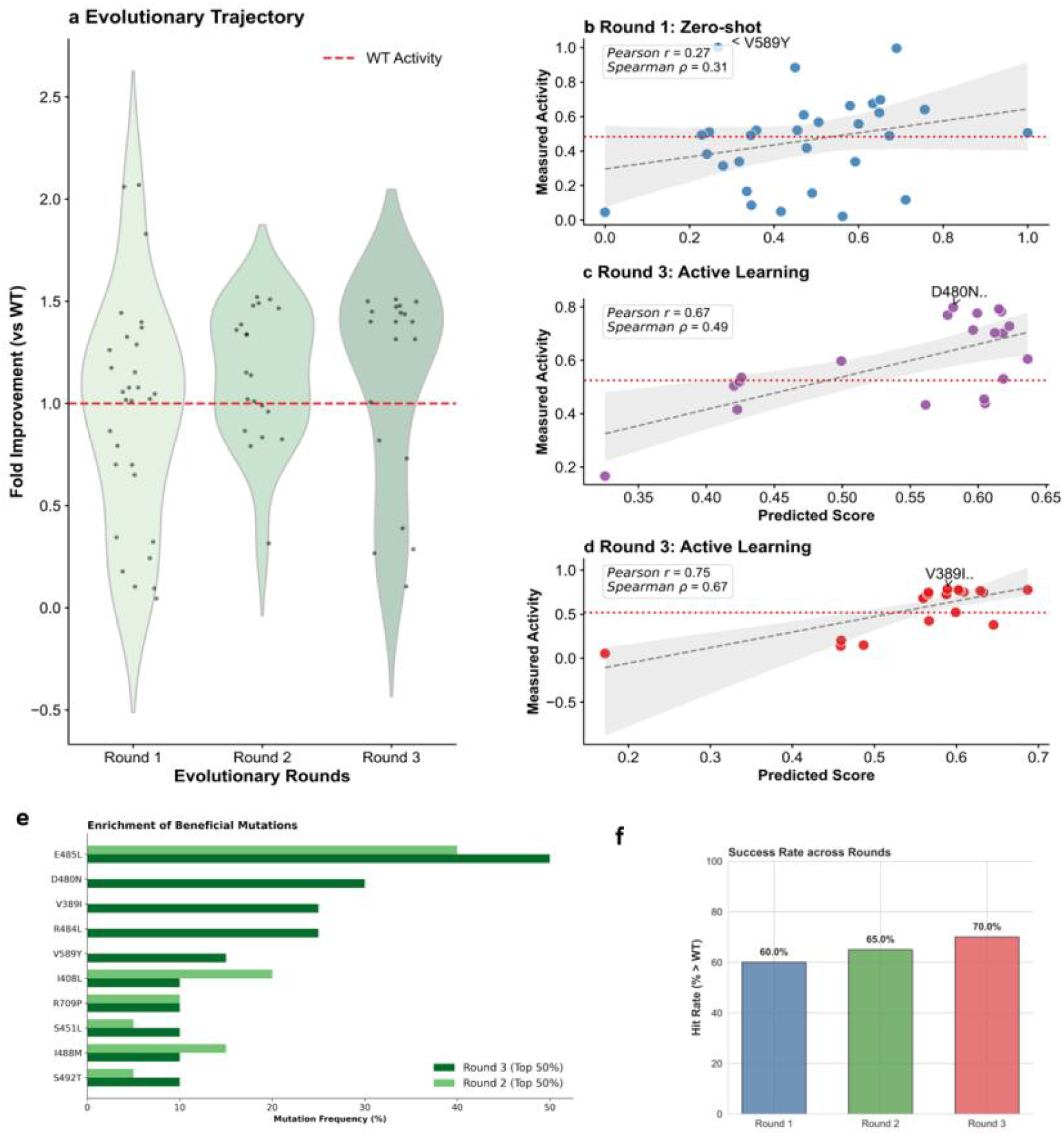
Evolutionary trajectory and model performance across iterative engineering rounds. (a) Evolutionary trajectory showing the fold improvement of variants over the wild-type (WT) enzyme across three rounds of screening. Round 1 (blue) utilized a zero-shot prediction strategy to identify initial hits from a broad sequence space, resulting in a substantial fold increase (up to ∼9.7-fold) over the low-activity parental WT. Round 2 (green) and Round 3 (dark green) employed the EVOLVEpro active learning framework, which successfully shifted the population distribution towards higher fitness. The red dashed line indicates the normalized activity of the WT control (Fold = 1.0). The violin plots depict the density of the activity distribution, with individual variants overlaid as black dots. (b– d) Correlation analysis between predicted fitness scores and experimentally measured activities for (b) Round 1 (Zero-shot), (c) Round 2 (Active Learning), and (d) Round 3 (Active Learning). The scatter plots demonstrate the model’s capability to rank variants, with the red dashed lines representing the internal WT activity for each specific round. Pearson (r) and Spearman (p) correlation coefficients are provided to quantify the predictive accuracy. Shaded areas represent the 95% confidence interval of the regression fit. (e) Experimental hit rates across the three iterative rounds, defined as the percentage of variants exhibiting catalytic activity superior to the corresponding wild-type (WT) or parental baseline. The zero-shot model (Round 1) achieved an initial hit rate of ∼60%. Notably, in the subsequent active learning rounds (Rounds 2 and 3), the hit rate maintained a high level (>66%) despite the increasing difficulty of improving upon optimized templates. (f) Mutational enrichment analysis comparing the frequency of specific amino acid substitutions within the top 50% of performing variants in Round 2 (light green) and Round 3 (dark green). The progressive enrichment of specific residues (e.g., I408L, E485L, R484L) in the final round provides molecular evidence that the EVOLVEpro framework effectively identifies and recombines epistatic hotspots to drive phenotypic improvement.

Zero-Shot Prioritization and the “Cold Start” (Round 1) In the initial exploration phase (Round 1), we faced the “cold start” challenge: identifying beneficial variants without prior experimental labels. We utilized a zero-shot protein language model (ESM-2) to prioritize single-point variants from a vast combinatorial space.

Performance: Despite the lack of training data, the zero-shot strategy successfully filtered out deleterious mutations, achieving an initial hit rate of ∼60.0% (percentage of variants superior to WT).

Best Hit: Notably, this broad sweep identified a standout variant with a 9.7-fold improvement in catalytic activity over the wild-type (WT) baseline (activity increased from 0.103 to 1.00 relative units).

Predictive Power: Correlation analysis revealed a positive trend (Pearson *r* = 0.27, Spearman *p* = 0.31) between the zero-shot likelihood scores and experimental activity. This confirmed the model’s intrinsic capacity to capture evolutionary constraints and identify functional hotspots even in a low-activity parental scaffold.

Defying Diminishing Returns with Active Learning (Rounds 2 & 3)

Building on the functional landscape mapped in Round 1, we transitioned to the EVOLVEpro active learning framework for subsequent rounds. By incorporating experimental feedback, the model was fine-tuned to capture the local fitness landscape and epistatic interactions.

Population Shift: As shown in the evolutionary trajectory (Fig. 4a), the population distribution shifted robustly towards higher fitness in Rounds 2 and 3. While the fold-improvement multiples appeared to stabilize due to the significantly elevated baseline of the starting templates, the absolute activity continued to climb.

Increasing Success Rate: Remarkably, the efficiency of our pipeline defied the “law of diminishing returns” typically observed in directed evolution. Instead of plateauing, the experimental hit rate increased from 60.0% (Round 1) to 65.0% (Round 2) and finally to 70.0% (Round 3).

Model Accuracy: This sustained success was underpinned by the model’s improving predictive accuracy. In Round 3, the correlation between predicted scores and measured activity strengthened significantly to Pearson *r* = 0.75 and Spearman *p* = 0.67. This demonstrates that the active learning loop effectively “learned” the genotype-phenotype map, enabling the precise identification of superior variants even within a highly optimized sequence space.

Molecular Mechanism: Epistatic Anchoring and Convergent Selection To elucidate the molecular basis of this rapid optimization, we analyzed the mutational enrichment across generations (Fig. 4f). The data reveals a distinct trend of convergent selection:

Enrichment: Specific beneficial substitutions identified in earlier rounds, such as I408L, E485L, and R484L, were significantly enriched in the top 50% of the Round 3 population compared to Round 2.

Key Mutations: Most notably, the frequency of the E485L mutation increased substantially (>50% frequency in Round 3), suggesting it serves as a critical “epistatic anchor”. Other mutations like D480N and V389I also showed high enrichment, indicating that the EVOLVEpro framework effectively learned to identify and recombine these synergistic building blocks to drive phenotypic improvement.

Collectively, these results demonstrate that our generative AI-driven pipeline can rapidly evolve enzymes from low-activity starting points to high-performance variants with minimal experimental screening (Low-N), validating the power of combining zero-shot priors with iterative active learning.

### 2.5 Benchmarking High-Fidelity Sequencing on the DNBSEQ Platform

To rigorously validate the translational potential of our AI-evolved enzymes in a commercial sequencing workflow, we utilized the automated BAL biofoundry to construct and purify the top-performing variants for head-to-head benchmarking. This final selection included DP1, a double mutant (E485L, R709P), and DP2, a highly complex variant containing five mutations (V389I, L453I, D480N, R484L, E485L). The successful automated construction of DP2, which required precise multi-site mutagenesis, serves as a testament to the robust “Build” capabilities of our platform.

We benchmarked these variants against a standard commercial polymerase (Reference) on the MGI DNBSEQ-E25 platform using the CoolMPS chemistry, processing an E. coli genomic DNA library (SE40). The sequencing run statistics revealed that the AI-engineered variants significantly outperformed the commercial standard across multiple critical quality metrics (Fig. 5).

**Figure 5.**
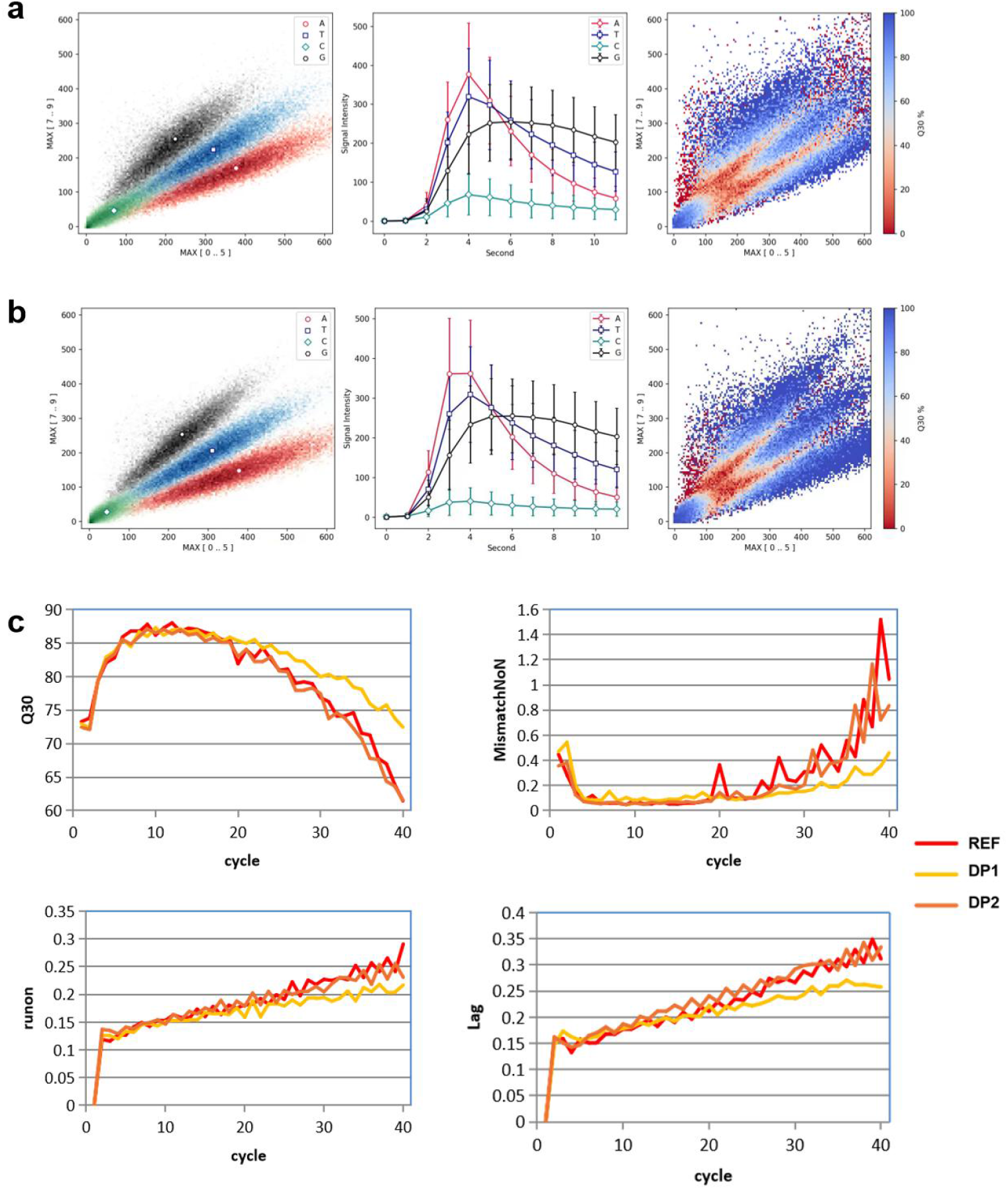
Benchmarking of AI-evolved polymerase variants on the DNBSEQ-E25 sequencing platform. Performance comparison of the engineered variants, DP1 (E485L, R709P) and DP2 (V389I, L453I, D480N, R484L, E485L), against the commercial reference polymerase (REF) using an E. coli genomic DNA library (SE40). (a) Summary table of key run statistics. The optimized variants demonstrated superior sequencing quality metrics, with DP1 achieving the highest Effective Spot Rate (ESR) and Q30 scores. (b) Analysis of signal stability and phasing kinetics. The engineered variants exhibited markedly reduced Lag (incomplete extension) and Runon (pre-phasing) rates compared to REF. This indicates that the AI-selected mutations improve the enzyme’s catalytic efficiency towards 3’-blocked nucleotides, leading to slower signal decay and higher data validity. (c) Error rate analysis showing a significant reduction in average base calling errors. DP1 achieved a 37% reduction in error rate (0.17% vs 0.27%), validating the high fidelity of the evolved enzymes.

Superior Data Yield and Effective Spot Rate (ESR) A key indicator of enzyme robustness in DNBSEQ is the Effective Spot Rate (ESR), which reflects the fraction of DNA nanoballs (DNBs) that support active sequencing. The engineered variant DP1 achieved a remarkable ESR of 73.22%, representing a substantial improvement over the Reference (66.49%). This suggests that the AI-selected mutations enhance the enzyme’s solubility or stability under the rigorous conditions of the sequencing flow cell, resulting in a higher yield of high-quality data.

Enhanced Fidelity and Reduced Phasing Errors In terms of sequencing accuracy, the engineered variants demonstrated exceptional fidelity.

Error Rate Reduction: The average error rate for DP1 was reduced to 0.17%, a 37% decrease compared to the Reference (0.27%). This superiority was corroborated by the Q30 scores, with DP1 reaching 90.02% versus 89.14% for the Reference.

Phasing Control (Lag/Runon): Crucially, the engineered variants exhibited superior control over phasing errors. The Lag rate (incomplete extension) for DP1 dropped to 0.31% (vs. 0.51% for REF), and the Runon rate (over-extension) decreased to 0.23% (vs. 0.36% for REF).

Mechanistic Implications of Evolved Mutations The significant reduction in Lag rates indicates that the engineered mutations — particularly E485L (present in both DP1 and DP2) and the supporting mutations in the complex DP2 variant — likely reshape the active site or the exit channel. This structural optimization facilitates the accommodation of the bulky 3’-blocked reversible terminators used in CoolMPS, ensuring more complete nucleotide incorporation in every cycle. This enhanced processivity translates directly into greater signal stability and slower signal decay over the sequencing run.

Ultimately, the best-performing variant yielded a mapping rate of 99.89%. These results not only confirm that our autonomous platform can evolve enzymes superior to current industrial standards but also demonstrate its capacity to navigate complex fitness landscapes to identify synergistic mutations (as seen in the 5-point mutant DP2) that are inaccessible to traditional screening methods.

**Table 1.** Sequencing performance of DP1 and DP2 variants on the DNBSEQ-E25 sequencer.

| Type | REF | DP1 | DP2 |
| --- | --- | --- | --- |
| Reference | Ecoli | Ecoli | Ecoli |
| CycleNumber | 40 | 40 | 40 |
| TotalReads(M) | 22.89 | 24.54 | 22.02 |
| ActivePixels(M) | 37.06 | 36.43 | 37.17 |
| LoadingRate(%) | 92.9 | 91.98 | 91.34 |
| MappedReads(M) | 22.84 | 24.51 | 21.98 |
| Q30(%) | **89.14** | **90.02** | **89.18** |
| Runon(%) | **0.36** | **0.23** | **0.31** |
| Lag(%) | **0.51** | **0.31** | **0.49** |
| ESR(%) | 66.49 | 73.22 | 64.86 |
| NonSpecificAdsorption(%) | 0.23 | 0.17 | 0.12 |
| MappingRate(%) | 99.78 | 99.89 | 99.82 |
| AvgErrorRate(%) | 0.27 | 0.17 | 0.24 |
| AvgErrorRate!N(%) | 0.27 | 0.17 | 0.24 |

## Discussion

Redefining Laboratory Automation via an AI-Native Architecture The core innovation of this study extends beyond the evolution of a specific polymerase; it lies in the establishment of a scalable, AI-native automated laboratory execution management system (Fig. 6). Traditional laboratory automation often suffers from “automation silos,” relying on rigid, device-specific scripts that are fragile and difficult to scale. In contrast, our platform introduces a “Cloud-Edge Synergistic” architecture that fundamentally decouples high-level scientific intent from low-level hardware execution. By deploying a real-time Edge Execution Layer to handle hardware abstraction (via RESTful/MCP protocols) and a Cloud Management Plane for global orchestration, we ensure both the reliability of local instrument control and the intelligence of global task scheduling. This architecture transforms the laboratory from a collection of isolated instruments into a logically unified, programmable digital entity.

Agent-Native Integration and Semantic Interoperability A distinct breakthrough of our system is its Agent-Native design. By deeply integrating the Model Context Protocol (MCP), we have bridged the semantic gap between Large Language Models (LLMs) and physical hardware. Unlike previous “human-in-the-loop” systems, our LLM agent can directly “perceive” instrument states and “actuate” physical workflows through standardized interfaces. This capability allows the system to autonomously parse natural language instructions (e.g., “optimize for lower error rate”) into structured JSON task graphs, automatically handling complex resource constraints and risk assessment. This represents a leap towards true “Lights-out Factories” for biotechnology, where dry-lab computation and wet-lab experimentation form a seamless, self-driving closed loop.

Overcoming Diminishing Returns with Active Learning Enabled by this robust infrastructure, our evolutionary campaign defied the “law of diminishing returns.” In classical directed evolution, identifying beneficial mutations becomes exponentially harder as the enzyme approaches an optimum. However, our EVOLVEpro-guided active learning strategy maintained a hit rate of >66% even in later rounds. This sustained efficiency suggests that the model successfully captured high-order epistatic interactions—such as the synergy between V389I and E485L—that act as “epistatic anchors” to stabilize the active site for non-natural substrates. The success of the CoolMPS polymerase serves as a validation that our AI-native infrastructure can effectively navigate rugged fitness landscapes that are inaccessible to manual or purely stochastic methods.

Standardization of the Physical-Digital Interface Finally, our system addresses the challenge of hardware heterogeneity through a “Meta-Action” protocol model. By abstracting device capabilities into brand-agnostic interfaces (e.g., a generic “Transfer Liquid” command rather than a specific Hamilton or Tecan instruction), we achieve zero-intrusive integration. This “Software-Defined Topology” allows for plug-and-play device addition and flexible reconfiguration of the control matrix without system downtime. As biofoundries scale, this ability to unify heterogeneous assets — software models, databases, and robotic arms — into a secure, traceable ecosystem will be critical for standardizing the development of industrial enzymes and therapeutic proteins.

## Methods

### System Software Implementation

The automated laboratory management system was built on a distributed microservices architecture implemented in Python 3.9 using the FastAPI framework.

Infrastructure: Services were containerized via Docker and orchestrated by Kubernetes (K8s) on a private cloud cluster. Data persistence relied on PostgreSQL for relational metadata and MinIO for unstructured object storage (Data Lake).

Edge Control: Edge nodes communicated with the cloud plane via gRPC (for synchronous commands) and RabbitMQ (for asynchronous telemetry).

Agent & MCP: The AI agent was powered by Qwen3-72B-Instruct served via vLLM. We implemented the Model Context Protocol (MCP) to expose the Hardware Abstraction Layer (HAL) as executable tools. The Agent generated Directed Acyclic Graphs (DAGs), which were validated by Pydantic schema and simulated in a Unity 3D digital twin environment before physical execution.

### Materials and Instrumentation

Oligonucleotides were synthesized by Sangon Biotech (Shanghai, China). Reagents for molecular biology included Phanta Flash Super-Fidelity DNA Polymerase (Vazyme, Suzhou, China), QuickCut ™ EcoRI (Takara, Beijing, China), and 10 × BugBuster Protein Extraction Reagent (Sigma-Aldrich, St. Louis, MO, USA). 3’-azidomethyl-modified nucleotides (3’-N3-dNTPs) were custom-manufactured by MGI (Wuhan, China). Purification consumables including Ni-NTA resin and Amicon Ultra centrifugal filters (MWCO 10 kDa/30 kDa) were obtained from Sangon and Sigma-Aldrich, respectively. Automated experimentation was executed on an MGISP-Smart8 liquid handling workstation (MGI, Shenzhen, China) integrated with a Cytomat 2 automated incubator (Thermo Fisher, Waltham, MA, USA). Microbial colony picking was performed using a QPix 400 system (Molecular Devices, San Jose, CA, USA), and fluorescence measurements were acquired on a Tecan Infinite 200 PRO microplate reader (Tecan, Männedorf, Switzerland).

### AI-Orchestrated Computational Framework

The autonomous workflow was coordinated by a central AI agent based on the Qwen3 large language model (LLM), accessed via a custom API interface to parse natural language requests into executable biofoundry protocols [31].

- Zero-Shot Prediction: Initial variant fitness was predicted using ESM-2 (esm2_t36_3B_UR50D), a transformer-based protein language model [14]. Masked language modeling (MLM) scores were calculated to rank variants based on the log-likelihood ratio relative to the wild-type sequence [32].
- Active Learning: For iterative optimization, we employed EVOLVEpro, a supervised regression framework designed for low-N protein engineering [23]. The model utilized frozen ESM-2 embeddings as input features and was updated between rounds using experimental data to refine the fitness landscape prediction [20].
- Phylogenetic Mining and In Silico ScreeningFamily B DNA polymerase sequences were retrieved from the InterPro database [33], filtered for “Thermophilic Archaea” taxonomy. Redundancy was reduced using CD-HIT with a sequence identity threshold of 90% [34]. The SoluProt predictor was subsequently applied to remove aggregation-prone sequences [35].

### Automated Library Construction and High-Throughput Screening

The gene encoding polyB was codon-optimized and synthesized by Azenta Life Sciences, then cloned into the pET-22b(+) vector and transformed into E. coli BL21(DE3).

- **Automated Mutagenesis**: Mutagenic primers were designed using the Vazyme CE Design tool. PCR mutagenesis was orchestrated by the MGISP-Smart8 system in 50 μL reactions containing 1 U Phanta Flash Polymerase, 0.2 ng/μL pET22b(+)-polyB template, and 0.4 μM primers. Following thermal cycling, reaction plates were unsealed and subjected to DpnI digestion (1 μL enzyme per 50 μL reaction) for 30 min at 37°C. The digested products were chemically transformed into BL21(DE3) competent cells using on-deck temperature control modules.
- **Expression and Lysis**: Single colonies were picked by QPix into 96-well plates containing LB medium (100 μg/mL ampicillin) and cultured overnight in the Cytomat 2. Cultures were subcultured (1:100) and grown at 37°C until log phase (OD600=0.6−0.8). Protein expression was induced with 1 mM IPTG for 4 hours at 37°C. Cell lysis was performed by adding 3 μL of 10× BugBuster to 150 μL culture, followed by 20 min incubation at room temperature.
- **Fluorescence-Coupled Activity Assay**: Enzymatic activity was quantified using a fluorescence-based coupled assay on the Tecan Infinite 200 PRO. The 50 μL reaction mixture contained 0.1 μM annealed DNA substrate (T28/P14/P13), 1× polyB reaction buffer (20 mM Tris-HCl pH 8.8, 10 mM KCl, 10 mM (NH4)2SO4, 20 mM MgSO4, 0.1% Triton X-100), 1× QuickCut Buffer, 2.5 μM 3’-N3-dTTP, 5 μL QuickCut™ EcoRI, and 10 μL crude lysate. Reactions were monitored at 40°C with fluorescence excitation/emission wavelengths of 490/525 nm at 80-second intervals. All reagent transfers were automated by the MGISP-Smart8 robotic arm.

### Gel-Based Validation of Crude Extracts

For qualitative validation, a 10 μL reaction system was employed containing 1 *μM* primer/template duplex (T40/P20), 1× polyB reaction buffer, 20 μM 3’-N3-dATP, and 1.5 μL crude lysate. Reactions were incubated at 40°C or 60°C for 10 min and quenched with an equal volume of 2×TBE-urea sample buffer. Products were resolved by 20% denaturing polyacrylamide gel electrophoresis (PAGE).

### Protein Purification and Characterization

Selected polyB variants were expressed in larger volumes as described above. Cells were harvested and resuspended in Lysis Buffer 50mM Tris−HCl, 500mMNaCl, pH7.0, followed by disruption via high-pressure homogenization. The supernatant was clarified by centrifugation and purified using Ni-NTA affinity chromatography. Eluted proteins were concentrated and buffer-exchanged into Storage Buffer 25 mM Tris−HCl, 150mM NaCl, pH7.0 using Amicon Ultra centrifugal filters (MWCO 30 kDa). Purified proteins were stored at -80°C. Specific activity was verified in a 10 μL reaction containing 0.05 mg/mL purified enzyme, 1 μM substrate (T18/P15), and 1 μM 3’-N3-dNTPs. Reactions were conducted at 60°C for 1 min and analyzed by denaturing 20% PAGE.

### Structural Modeling and Molecular Dynamics (MD)

To rigorously assess substrate stability, we employed a physics-based filtering pipeline:Structure Prediction:

- High-confidence 3D structures of the ternary complex (Polymerase-DNA-dATP) were generated using the Chai-1 folding algorithm, which has demonstrated superior accuracy for protein-ligand complexes [36].
- System Setup: MD simulations were performed using the Amber22 software package [37]. The protein and DNA were parameterized using the ff19SB force field [38], while the dATP substrate and modified nucleotides were parameterized using the General Amber Force Field (GAFF2) [39]. The system was solvated in an octahedral box using the OPC (Optimal Point Charge) water model [40], which provides improved accuracy for describing intrinsically disordered regions and hydration shells compared to TIP3P.
- Simulation Protocol: The system was neutralized with Na+ and Cl-ions. Following energy minimization, the system underwent gradual heating to 300 K and equilibration (50 ps NVT, 50 ps NPT). Production runs of 200 ns were executed at 300 K and 1 bar pressure [41].Energy Calculation: Binding free energies were calculated using the MM/GBSA (Molecular Mechanics/Generalized Born Surface Area) method [42]. Trajectories were sampled every 10 ps, and candidates with binding free energy < -40 kcal/mol were retained for experimental validation.

### Automated experimental screening workflow

All molecular biology operations were executed on an MGISP-Smart8 automated liquid handling workstation (MGI Tech) [43].

- Library Construction: Site-directed mutagenesis was performed via PCR using Phanta Flash Super-Fidelity DNA Polymerase (Vazyme). Plasmids were assembled using the Gibson Assembly method [44] and transformed into chemically competent E. coli BL21(DE3) cells.
- High-Throughput Expression: Cultures were grown in 96-deep-well plates, and protein expression was induced with 1 mM IPTG. Cells were harvested and lysed using BugBuster reagent (Millipore) [45].

### Functional Screening and Kinetic Characterization

Activity was monitored using a fluorescence-based incorporation assay on a Tecan Infinite 200 PRO microplate reader. The reaction mixture contained 3’-azidomethyl-modified nucleotides (3’-O-N3-dNTPs). Signal generation relied on the release of pyrophosphate or direct intercalation of SYBR Green I into the extended DNA product [46]. Kinetic parameters (Vmax, Km) were determined by fitting initial rates to the Michaelis-Menten equation.

### CoolMPS Sequencing Benchmarking

To evaluate the sequencing performance of the engineered polymerase variants in a realistic NGS workflow, benchmarking was performed using the CoolMPS chemistry on the DNBSEQ-E25 platform (MGI Tech)[24].

Library Construction and DNB Generation:

Genomic DNA from E. coli (strain MG1655) was fragmented and prepared using the MGIEasy DNA Library Prep Kit. The linear library molecules were circularized and amplified via Rolling

Circle Amplification (RCA) to generate DNA Nanoballs (DNBs) [47]. This amplification strategy avoids PCR-induced error propagation and ensures a high signal-to-noise ratio for downstream imaging. The concentration of DNBs was quantified using the Qubit ssDNA Assay Kit (Thermo Fisher) to optimize loading density [48].

Flow Cell Loading and Sequencing Configuration:

DNBs were immobilized onto patterned array flow cells to ensure uniform spacing and prevent optical overcrowding. For loading, 99 μL of the prepared DNB library was mixed with 33 μL of DNB Loading Buffer II, homogenized, and injected into the flow cell via the hydrodynamic loading port. Sequencing was executed in a Single-End 40 bp (SE40) configuration.

CoolMPS Biochemistry Cycle:

Unlike traditional sequencing-by-synthesis (SBS) methods that use fluorophore-labeled nucleotides, this study employed CoolMPS chemistry which relies on nucleobase-specific antibodies. The sequencing cycle consisted of:

1. Extension: The engineered polymerase incorporated unlabeled 3’-O-azidomethyl modified nucleotides (3’-blocked reversible terminators) into the growing primer strand.
2. Recognition: After washing, fluorescently labeled antibodies specific to the incorporated base were introduced to bind the generated epitope.
3. Imaging: Bioluminescent signals were captured by the DNBSEQ-E25 optical sensors.
4. Regeneration: The antibodies were displaced, and the 3’-blocking group was cleaved to permit the next cycle of extension.

Data Processing and Metrics:

Raw optical data were processed using the instrument’s built-in computing module for base calling. The resulting reads were aligned to the E. coli reference genome using the standard MGI analysis pipeline. To rigorously assess polymerase performance, the following quality metrics were calculated:

Effective Spot Rate (ESR): The percentage of array spots generating high-quality sequencing data, reflecting the efficiency of DNB loading and signal integrity.Q30: The percentage of bases with a quality score ≥ 30 (error probability ≤0.1).

Lag and Runon: Phasing metrics representing incomplete extension (Lag) or carry-over extension (Runon). These metrics serve as direct indicators of the polymerase’s processivity and catalytic efficiency in the context of the cPAS (combinatorial Probe-Anchor Synthesis) technology.

## Code availability

The code is currently being prepared for public release and will be available at Github shortly.

## Data availability

All data can be found in the supplementary materials.

## Author contribution

M. Y. conceived and designed the study. †C.W,Z, L.X.Y. contributed equally. C.W.Z., L.X.Y., Y.J.Q., and D.J.L. performed the analyses and drafted the manuscript with input from all co-authors. C.W.Z. and S.M.D. were responsible for the design, implementation, and operational management of laboratory automation in the biology laboratory. Y.J.Q. and D.J.L. contributed to the design and supervision of the experimental work. M.Y. provided critical guidance on manuscript writing and revision. All authors read and approved the final version of the manuscript.

## Acknowledgement

This research is supported by the Ministry of Science and Technology of the People’s Republic of China’s programme titled ‘National Key Research and Development Program of China’ (2022YFF1202200).

## Notes

### Competing Interest Statement

The authors have declared no competing interest.

